# Trans-omics analyses revealed differences in hormonal and nutritional status between wild and cultured female Japanese eel (*Anguilla japonica*)

**DOI:** 10.1101/483834

**Authors:** Masato Higuchi, Miyuki Mekuchi, Takeshi Hano, Hitoshi Imaizumi

## Abstract

Long-term stock decline in the Japanese eel (*Anguilla japonica*) is a serious issue. To reduce natural resource utilization in Japan, artificial hormonal induction of maturation and fertilization in the Japanese eel has been intensively studied. Recent experiment on feminized (by feeding a commercial diet containing estradiol-17β for first half year) cultured eels have shown ovulation problem, which is seldom observed in captured wild eels. In this study, we tried to investigate causes of ovulation problem frequently seen in cultured eels by comparative trans-omics analyses.

The omics data showed low growth hormone and luteinizing hormone transcription levels in the brain and low sex hormone–binding globulin transcription levels in the liver of the cultured eels. In addition, we found high accumulation of glucose-6-phosphate and, maltose in the cultured eel liver. We also found that docosahexaenoic (DHA) acid, eicosapentaenoic acid (EPA) and arachidonic acid (ARA) ratios in cultured eels were quite different from wild eels.

The data suggested that ovulation problem is due to prolonged intake of a high-carbohydrate diet and/or unbalanced DHA/EPA/ARA ratios in diet.

## Introduction

The Japanese eel (*Anguilla japonica*) is an important species for inland aquaculture in Japan because of its high economic value. The seed used in its cultivation is taken from wild glass eels captured from estuaries. However, as the glass eel arrival has been drastically dropped since 1970s, eel seed depletion has become a serious problem not only in Japan but also in other East Asian countries. To reduce natural resource utilization in Japan, artificial breeding techniques of eels have been studied, and rearing methods of eel leptocephali have been developed [1].

Cultured female and male eels do not mature sexually in captive conditions. Therefore, weekly injection of salmon pituitary extract (SPE) is administrated to captured wild female eels, acclimated to seawater, to induce oocyte maturation [2]. After 10–15 injections of SPE, 17α-hydroxyprogesterone is injected to induce final oocyte maturation and ovulation [3]. To induce maturation of male eels, recombinant eel luteinizing hormone is used (human chorionic gonadotropin was used in the past [4]).

The matured female and male eels are then transferred to a tank. When water temperature is raised from 20°C to 22°C, spawning takes place early in the morning. The Japanese eel has gymnovarian– type ovaries, which lack a part of the ovarian capsule; therefore, ovulated eggs are discharged into the abdominal cavity once and then spawned through the genital pore [5]. About 0.3–1.0 million fertilized eggs can be collected from one wild eel by using this method.

However, it has been observed that in feminized (by feeding a commercial diet containing estradiol-17β (E2) for the first half year) cultured eels, the genital pore is clogged with ovarian fragment during artificial spawning (Fig 1), therefore, normal release of eggs is severely disrupted.

**Fig 1.**
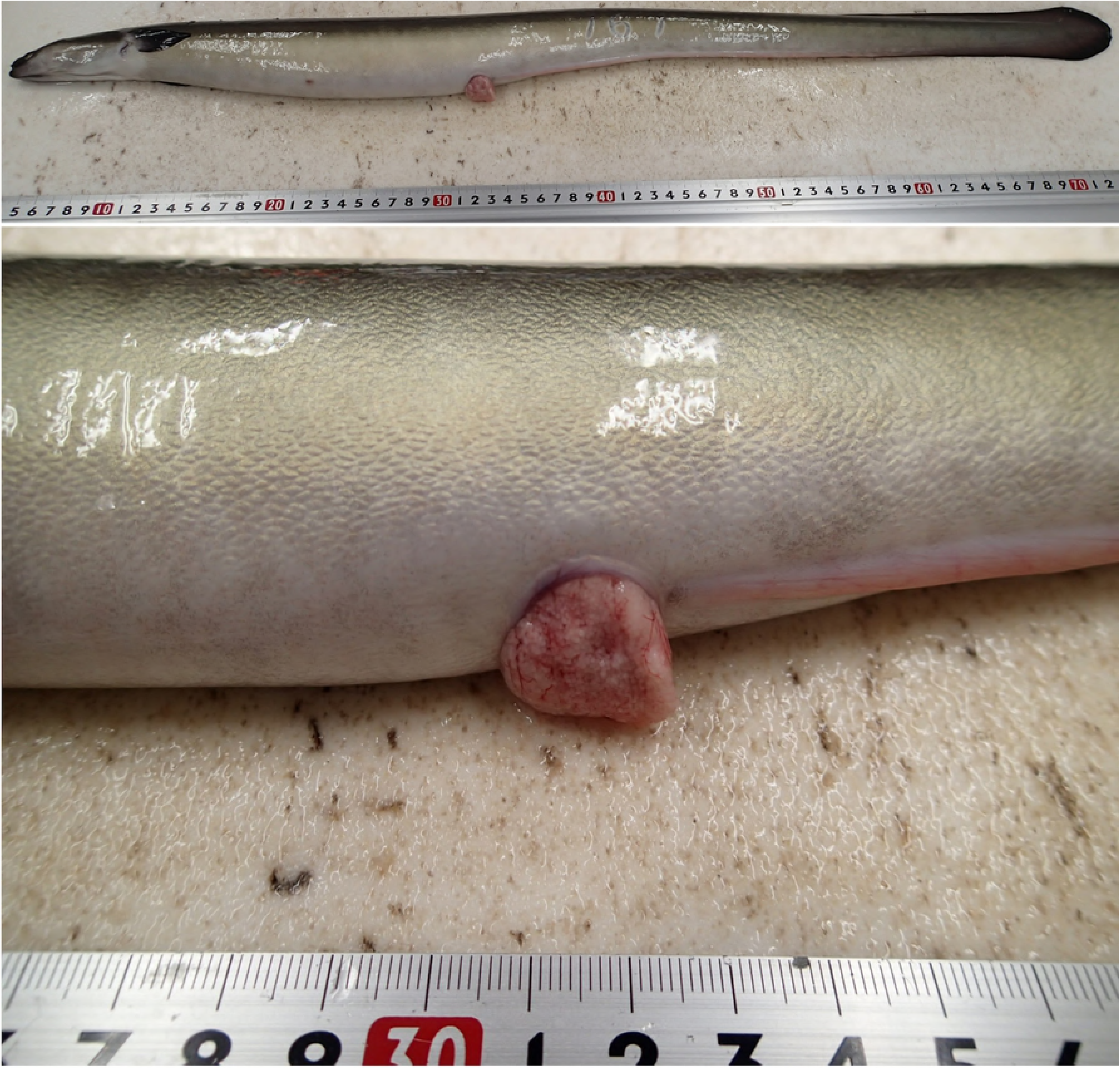
Clogged oviduct after hormone-induced spawning in cultured eel.

Anatomical observations have shown that many unovulated eggs remained in the ovary and ovarian fragment hypertrophy in cultured eels after artificial spawning, compared to fully ovulated eggs in wild eels. In 2015–2016, in 28.9 % (39/135) of cultured eels, the genital pore was clogged and more than 50-g eggs (approximately 2000 eggs/g) remained in the abdomen while in 1.6 % (1/61) of wild eel, the genital pore was clogged after artificial spawning. Furthermore, the average number of spawned eggs and the quality of fertilized eggs (e.g. fertilization, hatching, larval survival, and larval malformation rates) in cultured eels were lower than wild eels (data not shown). It seems that there is ovulation problem in cultured eels, but the reason is unclear. Hence, we hypothesized that the physiological status of the eels just before hormone injection was the reason for this difference since artificial induction of oocyte maturation method was common between cultured and wild eels.

In this study, we examined the differences between cultured and wild eels using combined transcriptomic and metabolomic analyses, also called trans-omics analysis. New omics technologies have already been applied to aquaculture study [6]. In comparison with wild eels, cultured eels showed quite different hormonal and nutritional status just before the beginning of induction of artificial oocyte maturation. These differences that will be clarified from our analyses would serve as keys to improving ovulation problems in cultured eels.

## Materials and Methods

### Ethics statement

All experiments were conducted in accordance with the institutional procedures of the National Research Institute of Aquaculture guidelines. The protocol was approved by the committee on the National Research Institute of Aquaculture (Permission No. 29016). All surgery performed under 2-phenoxyethanol anesthesia, and all efforts were made to minimize suffering.

### Animals

We purchased 20 wild female eels (from Lake Jinzai, Japan: 35^°^19^’^ N, 132^°^40^’^ E; body weight [BW]: 500–1000 g) from Mitani eel wholesaler, the local eel supplier in Shimane prefecture, on September 29, 2016. Acclimation to seawater was followed by usual procedure [7]. Briefly, the eels were stocked in a 500-L tank, into which seawater (26–27°C) was poured. After 36 h, the salinity was 33 ‰. Afterwards, 5 eels were randomly selected and transferred to a 2-kL semi-intensive tank filled with seawater, and the water temperature was gradually decreased from 26°C to 20°C (at the rate of 1.5°C /day). At the end of temperature adjustment, eels were then kept in the tank for 14 days without feeding and without light until the sampling date.

To obtain wild male eel, we purchased 20 small-size wild eels (BW: 180–250 g) from the same eel supplier. The eels were acclimated to seawater and water temperature of 20°C as mentioned earlier and reared for 14 days without feeding and without light until the sampling date. Gender was determined by morphological observation of the gonads. We found that 5 eels were male, and the rest were female.

For feminization experiment, we purchased 1000 juvenile grass eels from Oosumi fishery cooperative, the commercial eel supplier in Kagoshima prefecture on February 1, 2013. After acclimation to commercial fish feed, eels were fed on a diet with E2 (10 mg/kg diet) [8] every day until July 29, 2013, after which the eels were fed modified commercial feed without E2 supplementation (Table 1) on every Monday, Wednesday, and Friday. The feeding ratio was periodically adjusted on the basis of appetite. The eels were reared in fresh water at natural water temperature until September 29, 2016. Then 5 feminized cultured eels (BW: 800–900 g) were randomly selected and transferred into a 500-L tank and same acclimation treatments as in the above mentioned were carried out, and they were kept in the tanks for 14 days without feeding and without light until the sampling date.

**Table 1.**
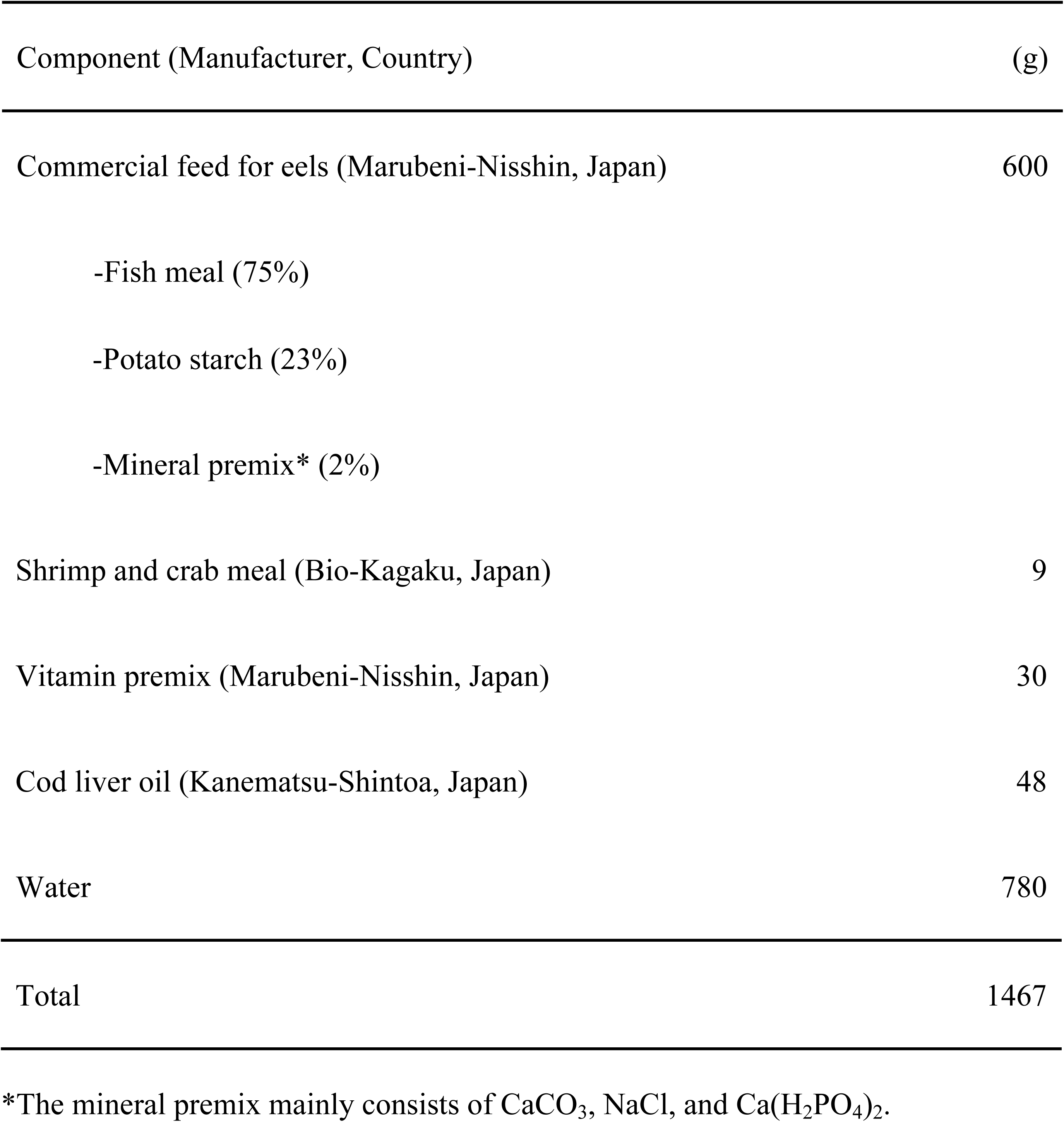
Diet formulation for cultured eels

### Tissue collection

The eels were anesthetized with 0.5% 2-phenoxyethanol. After total length and BW measurements, the brain, liver, ovaries, and blood were sampled from each eel; Table 2 shows the data collected. The whole brain and 5-mm^3^ pieces of the liver and ovaries were immediately placed in RNAlater solution (Ambion, Austin, TX, USA) and stored at −25°C. The liver and ovary samples were then weighed and sliced into approximately 10-mg pieces, frozen in liquid nitrogen, and stored at −80°C until analysis. We also fixed 5-mm^3^ pieces of the ovary samples in 10 % neutral phosphate-buffered formalin and stored at 4 °C until analysis. The remaining liver and ovary samples were stored at −80°C. The blood samples were stored for 3h at 4°C and then centrifuged at 1300 × g for 10 min at 4°C. Subsequently, the serum was collected, aliquoted, and then stored at −80°C until analysis.

**Table 2.** The collected data from cultured and wild eels at the day of sampling Values are expressed as the mean ± standard error of mean (SEM) of 5 samples Condition factor = 100 × weight(g)/length(cm)^3 Hepatosomatic index: HSI = [liver weight/body weight] × 100. Gonadosomatic index: GSI = [ovary weight/body weight] × 100. * *P < 0.05*.

|  | Cultured | Wild |
| --- | --- | --- |
| Total length (cm) | 80.8 ± 1.1 | 75.4 ± 1.8 |
| Body weight (g) | 841.8 ± 20.3 | 766.2 ± 45.5 |
| Condition factor | 0.159 ± 0.006 | 0.179 ± 0.005* |
| Liver weight (g) | 10.7 ± 1.2 | 8.5 ± 0.2 |
| HSI | 1.3 ± 0.1 | 1.1 ± 0.1 |
| Gonad weight (g) | 9.6 ± 0.9 | 12.4 ± 0.6* |
| GSI | 1.1 ± 0.1 | 1.6 ± 0.1* |

Values are expressed as the mean ± standard error of mean (SEM) of 5 samples
Condition factor = $100 \times \text{weight(g)}/\text{length(cm)}^3$
Hepatosomatic index: HSI = $[\text{liver weight}/\text{body weight}] \times 100$ .
Gonadosomatic index: GSI = $[\text{ovary weight}/\text{body weight}] \times 100$ .
\* $P < 0.05$ .

### Transcriptome analysis

Total RNA was extracted using the RNeasy Lipid Tissue Mini Kit (Qiagen, GmbH, Hilden, Germany) according to the manufacturer’s instruction. We evaluated the RNA quality of the whole brain and the liver on the basis of the proportion of ribosomal RNA (rRNA) in them using the Agilent 2100 Bioanalyzer Nano 6000 Kit (Agilent technologies, Palo Alto, CA, USA). The average RNA integrity numbers (RINs) of the whole brain and the liver were 9.0 and 7.0, respectively. However, since several reports have shown that due to unusual RNA composition, RINs of ovary samples in primary growth stages cannot be calculated using the RIN algorithm [9], we did not analyze the RINs of the ovary samples. In all, 4 total RNA (500 ng) samples (no.1 -no. 4) from each eel group (cultured and wild) were processed using the TruSeq RNA Sample Preparation v2 Kit (illimuna, San Diego, CA, USA) according to the manufacturer’s instructions. We constructed cDNA libraries with barcodes in order to enable multiplexing of pools, and sequenced the libraries using the illumina Hi-Seq 4000 platform equipped with a 50-base pair (bp) single-end module. The sequence data was deposited into DDBJ sequence read archive. The accession numbers are DRA007019 (brain), DRA007045 (liver), and DRA007046 (ovary), respectively. For postsequencing, raw reads were filtered by removing low-quality reads (quality value [QV] < 20). Next, we performed read-mapping analysis using CLC Genomics Workbench 8.1 (https://www.qiagenbioinformatics.com/) using the predicted protein-coding genes in the Japanese eel genome as a reference [10], and the expression value was calculated using the total number of reads mapped to the genes. Statistical analyses included the digital gene expression (DGE) test of CLC Genomics Workbench 8.1, and differences at *p <0.05* were considered statistically significant.

### Quantitative Polymerase Chain Reaction

After priming with a random hexamer, first-strand cDNA was synthesized from 1 μg of total RNA using the Omniscript RT Kit (Qiagen GmbH, Dusseldorf, Germany). Primers and probes were designed in reference to the Japanese eel genome data mentioned before [10]; see Supplemental table S1 for primer and probe sequences. The PCR mixture comprised each primer pair (0.5 μM) and probe (0.2 μM), and the FastStart Essential DNA Probes Master (2x) (Roche Diagnostics, Switzerland). The data was analyzed using the CFX96 Touch Real-time PCR Detection System (Bio-Rad, Hercules, CA, USA). The expression levels of messenger RNA (mRNA) in the samples were normalized to 60S acidic ribosomal protein P0 (*rplp0*) mRNA expression and calculated using the 2^−∆∆Ct^ method.

### Metabolomics analysism

The chemicals we used in the gas chromatography-mass spectrometry (GC-MS)-based metabolomics study were as follows: methoxyamine hydrochloride (Sigma Aldrich, St. Louis, MO, USA)); pyridine (Kanto Kagaku Chemical, Japan); pesticide-analytical-grade chloroform, HPLC-analytical-grade methanol, 2-propanol, acetonitrile, and ultrapure water (Wako Chemical, Japan); and N-methyl-N-(trimethylsilyl)-trifluoroacetamide (MSTFA) (GL Science Inc., Japan). Ribitol solution (Wako, Japan) in MilliQ water (200 mg/L) was used as an internal standard. Also, d27-Myristic acid (400 mg/L, Sigma Aldrich) was prepared in methanol in order to lock the retention time as described later.

To perform metabolomics on the liver and ovaries [11], we combined 1 mL of solvent mixture (CH3Cl:CH3OH:H2O, 1:2.5:1, v/v) and 60 µL of ribitol solution (10 mg ribitol dissolved in 100 mL water) with a tissue sample in a 2-mL microcentrifuge tube. Zirconia balls (1 mm in diameter) were added to the tube, and the sample was subjected to a sample crusher system (Fast Prep 24 Instrument, Funakoshi, Tokyo, Japan) for 1 min. The sample was then shaken for 30 min at 37°C and centrifuged at 16,000 × *g* for 5 min at 4°C. The supernatant (900 µL) was mixed with 400 µL of water and centrifuged again at 16,000 × *g* for 5 min, resulting in the formation of two phases: water phase (supernatant) and organic phase (bottom layer). The supernatant (800 µL) was mixed with 50 µL of d27-myristic acid solution and transferred to a 1.5-mL Eppendorf tube with a pierced cap. In addition, 100 µL of organic phase was collected and mixed with 50 µL of d27-myristic acid solution.

To prepare a serum, 50 µL of each sample were quenched by using 0.25 ml of a solvent mixture (acetonitrile:2-propanol: ultrapure water, 3:3:2, v/v) and mixed with 60 µL of ribitol and 50 µL of d27-myristic acid solution. The resulting extraction mixture of liver, ovaries, and serum was completely dried in a vacuum centrifuge drier (CVE-3000, Tokyo Rikakikai, Japan). For all samples, except the organic phase extracts of liver and ovaries, we performed two-step derivatization: (i) oximation and (ii) silylation. For oximation, we added 100 µL of methoxyamine hydrochloride in pyridine (20 mg/L) to the samples, and incubated the mixture for 90 min at 30°C. Thereafter, we added 50 µL of MSTFA to the samples for silylation, followed by incubation for 30 min at 37°C. The organic phase extracts were dissolved in 100 µL of pyridine followed by silylation with 50 µL of MSTA for 30 min at 37°C.

GC separation of the metabolites was performed using an Agilent 6890N gas chromatograph (Agilent technologies, Japan), equipped with a quadrupole mass spectrometer (Agilent 5975) and a DB-5ms column (i.d. 0.25 mm × 30 m, 0.25 μm thickness, 10 m DG, Agilent technologies). The samples (1 µL each) were injected in split mode (25:1, v/v); the temperatures of the injector, transfer line, and ion source were 250°C, 275°C, and 250°C, respectively. The helium gas flow rate was adjusted to 1.1 mL/min for a d27-myristic acid retention time of 16.727 min, and the oven temperature program was set as follows: 60°C for 10 min, increased at a rate of 15 °C/min to 325°C, and then held at 325°C for 3 min.

For data processing, we performed ion chromatographic deconvolution using the Automated Mass Spectral Deconvolution and Identification System (AMDIS, ver.2.7, Agilent, CA, USA). Metabolites were identified by their mass spectral patterns and retention times using the Fiehn Library (Agilent) as implemented in MSD ChemStation (Agilent). We successfully identified 61 metabolites (51 for the water phase, and 10 for the organic phase), 58 metabolites (47 for the water phase, and 11 for the organic phase) and 65 metabolites in the tissue samples of the liver, ovaries and serum, respectively.

### Serum lipids quantification

Serum cholesterol, triglyceride and phospholipids levels were measured using the Wako series test kit (Wako, Japan) according to the manufacturer’s microplate instructions.

### Serum and ovarian steroids quantification

Liquid chromatography-tandem mass spectrometry (LC-MS/MS) was used to assay serum estrone, E2, androstenedione, testosterone (T), 5α-dihydrotestosterone (5α-DHT), 11-ketotestosterone (11-KT), progesterone and 17-hydroxyprogesterone levels and ovarian E2 and T levels. For detailed procedures refer to other studies [12].

### TUNEL assay

Results of transcriptomic analysis of the ovary indicated that apoptosis occurs in cultured eel ovaries due to high expression of apoptosis-associated genes, such as small nuclear ribonucleoprotein F, tumor necrosis factor receptor superfamily member 19, cytochrome c, and granzyme K. Therefore, we used the terminal deoxynuculeotidyl transferase triphosphate (dUTP) nick end labeling (TUNEL) to confirm that cell death occurs through apoptosis. Ovary samples were fixed in 10% neutral phosphate-buffered formalin, embedded in paraffin wax and sectioned at 5-μm thickness. Then TUNEL assay was performed using ApopTag^®^ plus *in situ* apoptosis detection kit (Merck Millipore, UK) and hematoxylin counterstaining according to the manufacturer’s instructions.

## Statistical analysis

We used the *t*-test to measure statistical differences in the gonad somatic index (GSI), hepatosomatic index (HIS), serum lipids levels, serum steroid levels, ovarian steroid levels, and quantitative PCR analyses.

For qPCR of the brain samples of cultured female, wild female, and wild male, we determined statistical differences by one-way ANOVA, followed by Tukey’s test.

For metabolite analysis, for water phase (liver and ovary samples) and the serum, we normalized unique masses of each metabolite peak using the unique mass of the ribitol peak (m/z 217). For the organic phase of the liver and ovary samples, we performed normalization using the unique mass of the d-27 myristic acid peak (m/z 312). To identify key metabolites that help differentiate between cultured and wild eels, we performed orthogonal partial least-square discriminant analysis (OPLS-DA) followed by the S-plot using SIMCA 14.0 (Umetrics, Umea, Sweden). Pareto (Par) scaling of the normalized value was selected before fitting. MS-DIAL was used to find the differences in metabolites between two experimental eel groups using the *t*-tests for parametric data and Welch’s *t*-tests for nonparametric data at *p < 0.05* [13].

## Results

### Transcriptomic assembly

Transcriptomic analysis was carried out to evaluate the differences in gene expression patterns between wild and cultured eels. The cDNA libraries constructed from eel liver, ovary, and brain samples were sequenced using Illumina Hi-Seq 4000. From each individual sample, approximately 20 million sequence reads were obtained on average, and approximately 70% of the reads were mapped to the reference sequences. mRNA transcripts satisfied the *p < 0.05* criterion of the Gaussian-based test, and a fold change > 2.0. For further analysis, normalized expression values > 100 in the higher-expressed eel group were used (See Supplemental table S2–S5) for a list of screened mRNA transcripts.

### Gene expression comparison

On the basis of the results of RNA-seq (Supplemental table S2–S4), we performed validation and further analysis using real-time quantitative PCR (RT-qPCR). In the liver samples, we examined mRNA expression of 3-oxo-5-beta-steroid 4-dehydrogenase (*akr1d1*), cytochrome P450 1A9 (*cyp1a9*), and sex hormone-binding globulin (*shbg*) because these genes are related to steroidogenesis and steroid transport. We observed the mRNA expression level of *akr1d1* was 1.3-fold higher in cultured eels compared to wild eels, while *cyp1a9* and *shbg* mRNA levels were 8.9-fold and 6.6-fold higher, respectively, in wild eels compared to cultured eels (Fig 2).

**Fig 2.**
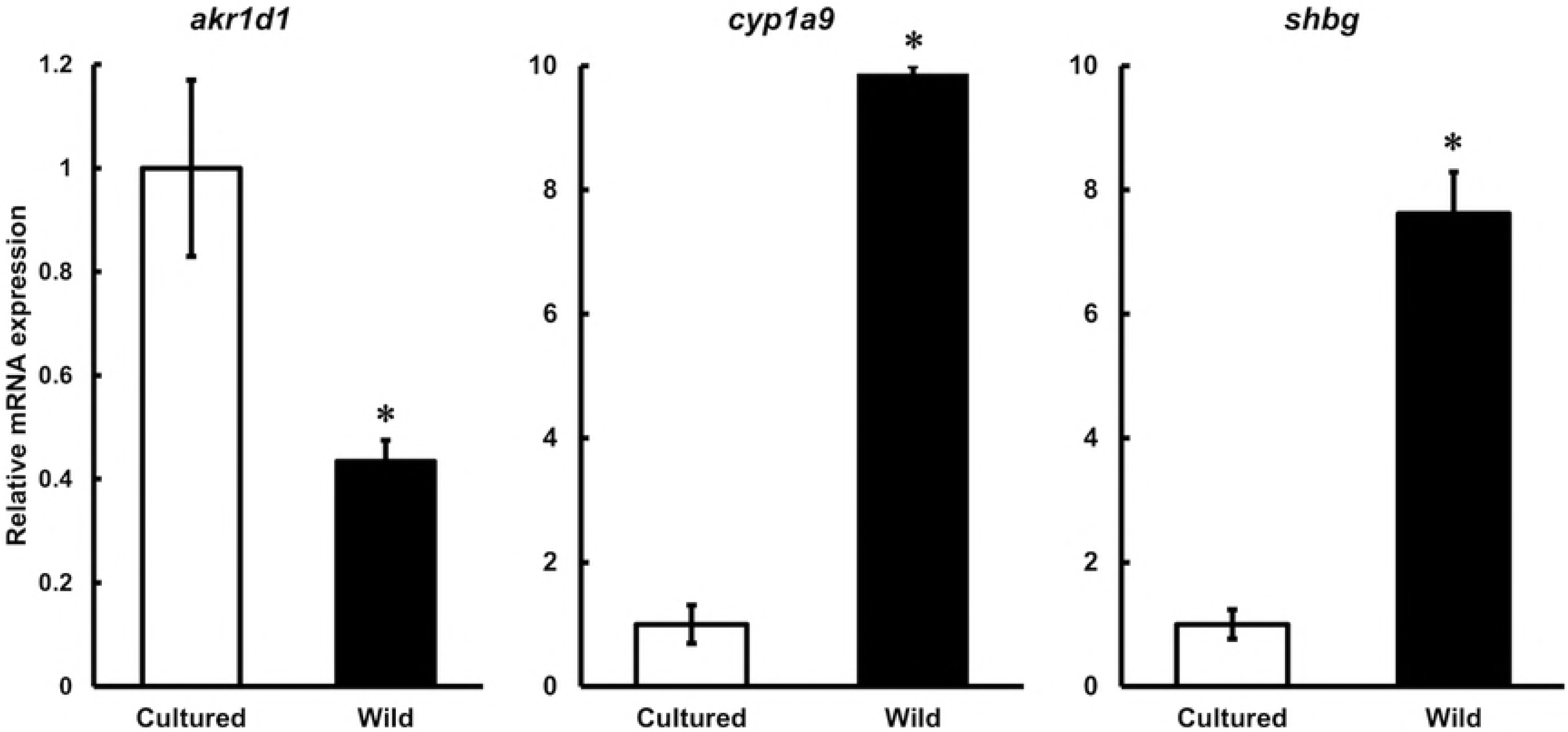
RT-qPCR analysis of liver samples. RT-qPCR analysis of 3-oxo-5-beta-steroid 4-dehydrogenase (*akr1d1)*, cytochrome P450 1A9 *(cyp1a9)*, and sex hormone-binding globulin (*shbg)* mRNA expression in cultured and wild eel liver samples Relative mRNA expression is normalized against 60S acidic ribosomal protein P0 *(rplp0)* mRNA expression. Each vertical bar represents the mean ± SEM of 5 samples. *\*P < 0.01*.

In the brain samples, we analyzed the expression levels of *follicle-stimulating hormone* (*fsh*), *luteinizing hormone* (*lh*), and *growth hormone* (*gh*) because these peptide hormones are closely related to reproduction. Although no significant difference was observed in the *fsh* transcription levels between wild and cultured eel (same as in the case of RNA-seq), the *lh* and *gh* transcription levels in wild eels were 1.5-fold and 4.5-fold higher, respectively, than in cultured eels (Fig 3A).

**Fig 3.**
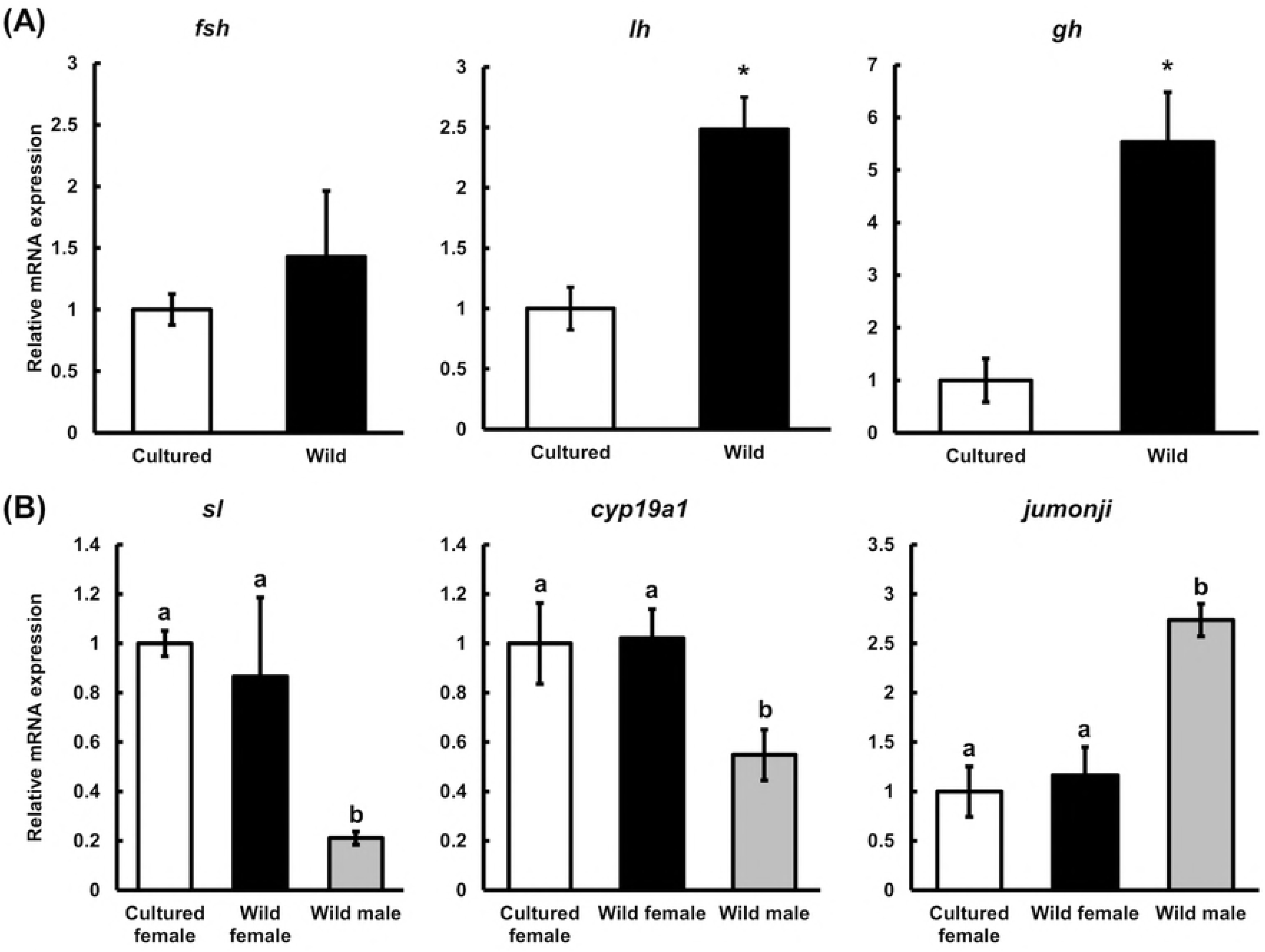
RT-qPCR analysis of eel brain samples. (A) RT-qPCR analysis of follicle stimulating hormone *(fsh)*, luteinizing hormone *(lh)*, and growth hormone (*gh)* in cultured and wild eel brain samples. Relative mRNA expression is normalized against 60S acidic ribosomal protein P0 (*rplp0*) mRNA expression. Each vertical bar represents the mean ± SEM of 5 samples. *\*P* < *0.05*. (B) RT-qPCR analysis of somatolactin (*sl)*, cytochrome P450 19A1 (*cyp19a1)*, and protein jumonji (*jumonji)* mRNA expression in the brain samples of cultured female, wild female, and wild male eels. Relative mRNA expression is normalized against 60S acidic ribosomal protein P0 (*rplp0*) mRNA expression. Each vertical bar represents the mean ± SEM of 5 samples. *a and b* indicate significant differences (*P*< 0.05).

In cultured eels, due to lower *shbg* transcription levels, we suspected hyperandrogenism and masculinization. To confirm this, marker genes showing sex-dependent expression patterns from Supplemental table S5. *somatolactin* (*sl*) and *cytochrome P450 19A1* (*cyp19a1*) were selected as female highly expressed genes, and *protein jumonji* (*jumonji*) was selected as the male highly expressed gene. No significant differences were observed between wild and cultured eels in the selected gene expression levels; however, there were significant differences between female and male eels (Fig 3B). Results of transcriptomic analysis showed that the expression levels of the four JUMONJI family genes are different between wild female and wild male in the brains. Amino acid sequences of four JUMONJI family genes contain ARID, jmjC and/or jmjN domains, which characterize JUMONJI; however, the sequences show some differences from one another. Primers and probe were designed for the no. 6 of in the Supplemental table S5 because only no. 6 contains all three ARID, jmjC, and jmjN domains. The mRNA expression pattern obtained from qPCR was the same as that obtained from RNA-seq data.

### Serum lipids and steroids, and ovarian steroid levels

Serum cholesterol, triglyceride, and phospholipids levels were measured (Table 3). There were no significant differences in cholesterol and phospholipid levels between wild and cultured eels, while serum triglyceride levels of cultured eels were significantly higher than those of wild eels.

**Table 3.** Serum lipid, steroid levels, and ovarian steroid levels in eels Values are expressed as the mean ± SEM (n=5). * *P < 0.05*.

|  | Cultured | Wild |
| --- | --- | --- |
| **Serum lipid levels** |  |  |
| Cholesterol (mg/dL) | 315.4 ± 35.1 | 303.4 ± 29.1 |
| Triglyceride (mg/dL) | 971.4 ± 145.2 | 543.0 ± 55.0* |
| Phospholipids (mg/dL) | 659.7 ± 58.3 | 731.2 ± 42.5 |
| **Serum steroid levels** |  |  |
| Estradiol-17β (pg/mL) | 99.5 ± 28.5 | 85.6 ± 26.6 |
| Androstenedione (pg/mL) | 12.1 ± 3.5 | 12.2 ± 4.8 |
| Testosterone (ng/mL) | 0.63 ± 0.2 | 1.03 ± 0.5 |
| 5α-Dihydrotestosterone (pg/mL) | 23.2 ± 4.1 | 37.8 ± 14.1 |
| 11-Ketotestosterone (ng/mL) | 0.64 ± 0.2 | 1.14 ± 0.2 |
| 17-Hydroxyprogesterone (pg/mL) | 76.1 ± 38.4 | 43.1 ± 6.2 |
| **Ovarian steroid levels** |  |  |
| Estradiol-17β (pg/g-wet) | 68.7 ± 17.9 | 62.1 ± 23.5 |
| Testosterone (pg/g-wet) | 129.4 ± 29.5 | 223.7 ± 77.3 |

LC-MS/MS was used to assay total serum estrone, E2, androstenedione, T, 5α-DHT, 11-KT, progesterone and 17-hydroxyprogesterone levels, and ovarian E2 and T levels (listed in Table 3). The estrone and progesterone levels were under the lower limit of quantitation (1 pg/mL and 2 pg/mL, respectively), and therefore were omitted. Serum T and 11-KT levels were lower in cultured eels than in wild eels, although no significant differences were observed.

Ovarian T levels were also lower in cultured eels, although, again, there were no significant differences.

### Metabolomics

For metabolomic analysis, we performed OPLS-DA followed by the S-plot in order to estimate the crucial metabolites contributing to differences between cultured and wild eels. For detailed procedures refer to other study [14]. For all tissues, the predictive component t[1] successfully separated the two eel groups. (Fig 4A, B, 5A, B, and 6A). Candidate biomarkers, with an absolute p(corr) value of greater than 0.6, were selected [15] (Fig 4C, D, 5C, D, and 6B). R2 and Q2 were used to evaluate the model. R2 represents the model’s goodness. R2X is the fraction of the variation of the X variables, and R2Y is the fraction of the variation of Y variables explained by the model. Q2 represents the model’s goodness of prediction and also an estimate of the model’s predictive ability. Q2 is calculated by cross-validation. The higher the values of R2 and Q2, the better is the model’s predictive ability.

**Fig 4.**
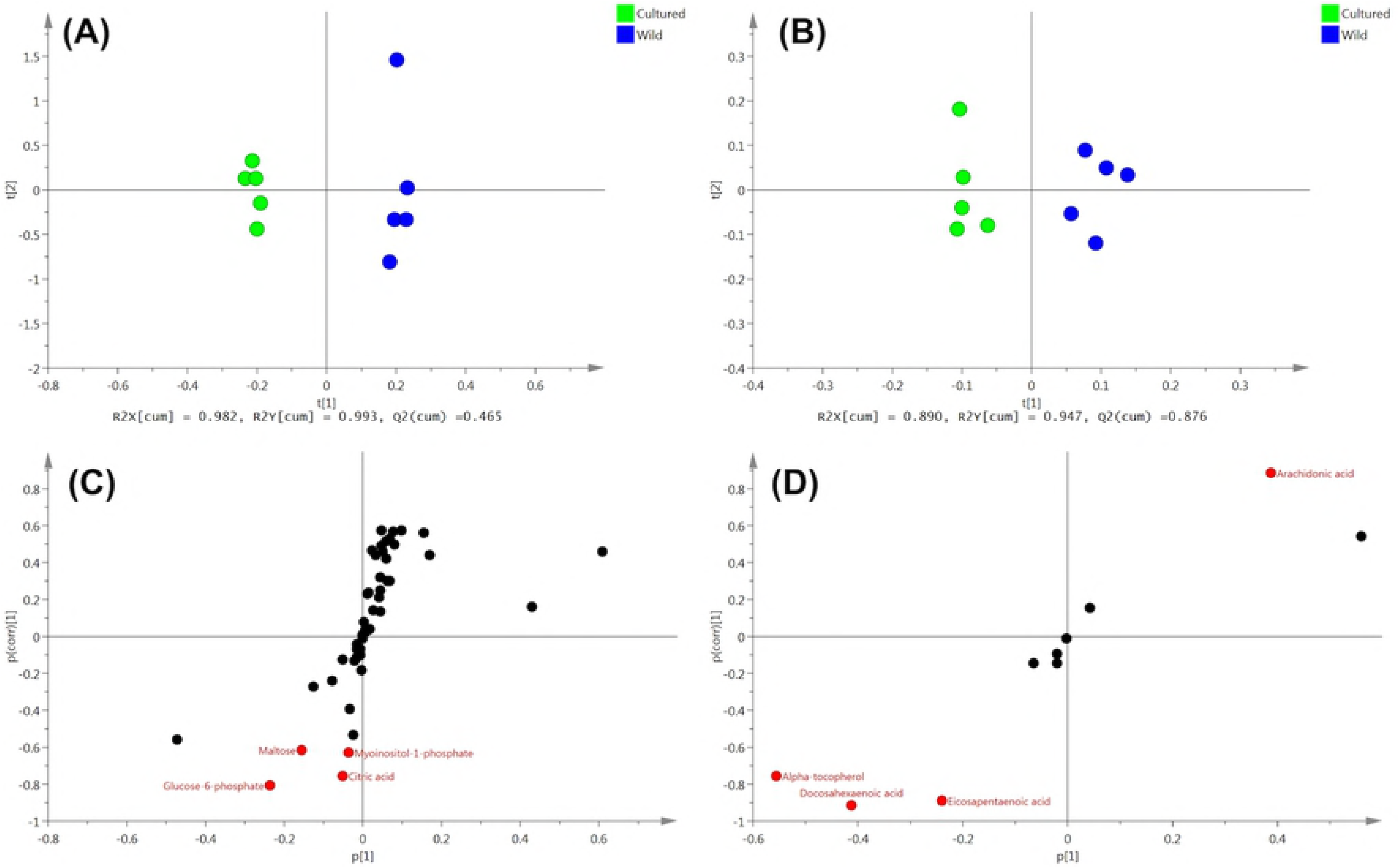
OPLS-DA of liver metabolites obtained from cultured and wild eels. (A) water phase fraction and (B) organic phase fraction. S-plot generated from OPLS-DA of (C) water phase fraction and (D) organic phase fraction.

**Fig 5.**
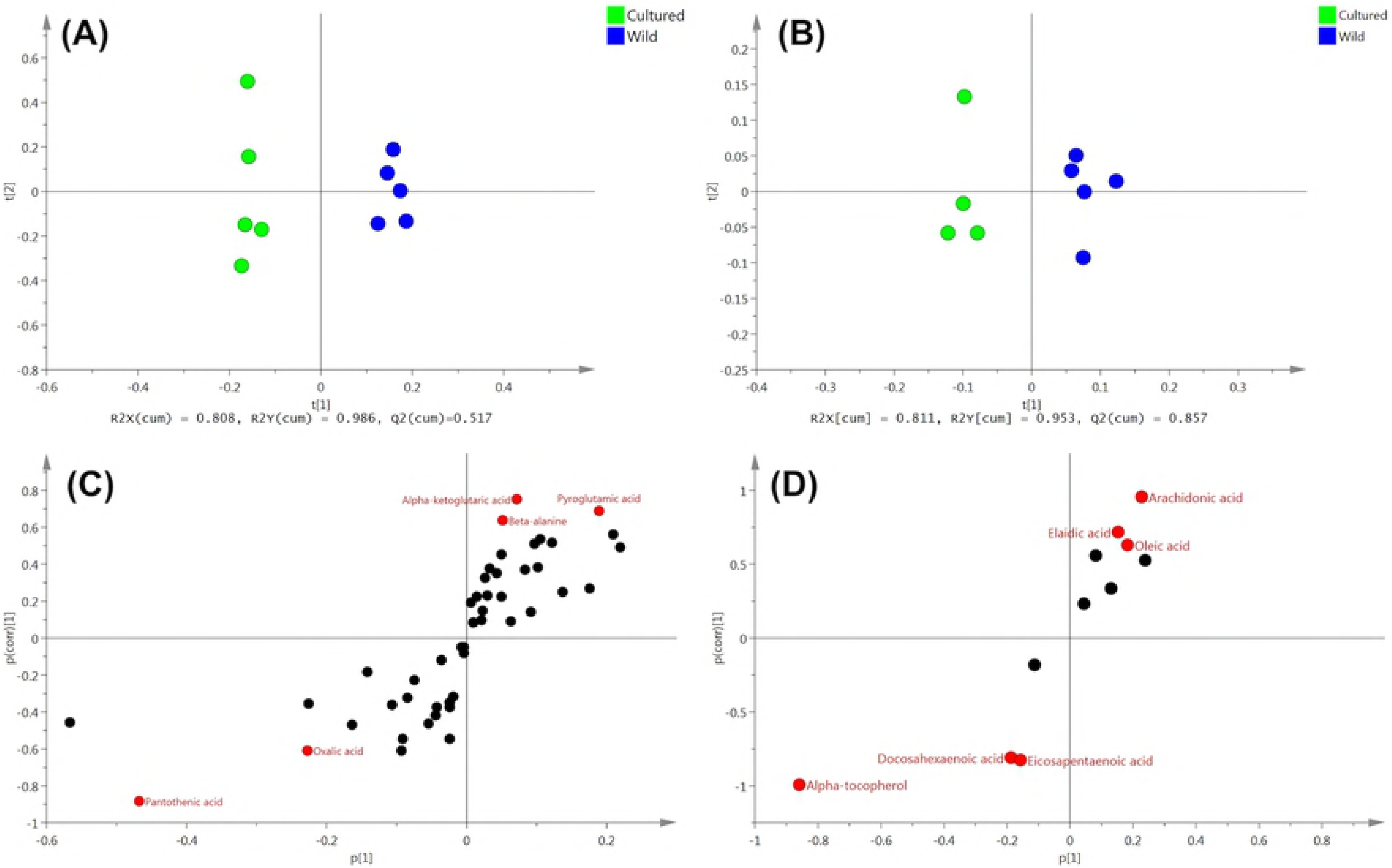
OPLS-DA of ovary metabolites obtained from cultured and wild eels. (A) water phase fraction. (B) organic phase fraction. S-plot generated from OPLS-DA of (C) water phase fraction and (D) organic phase fraction.

**Fig 6.**
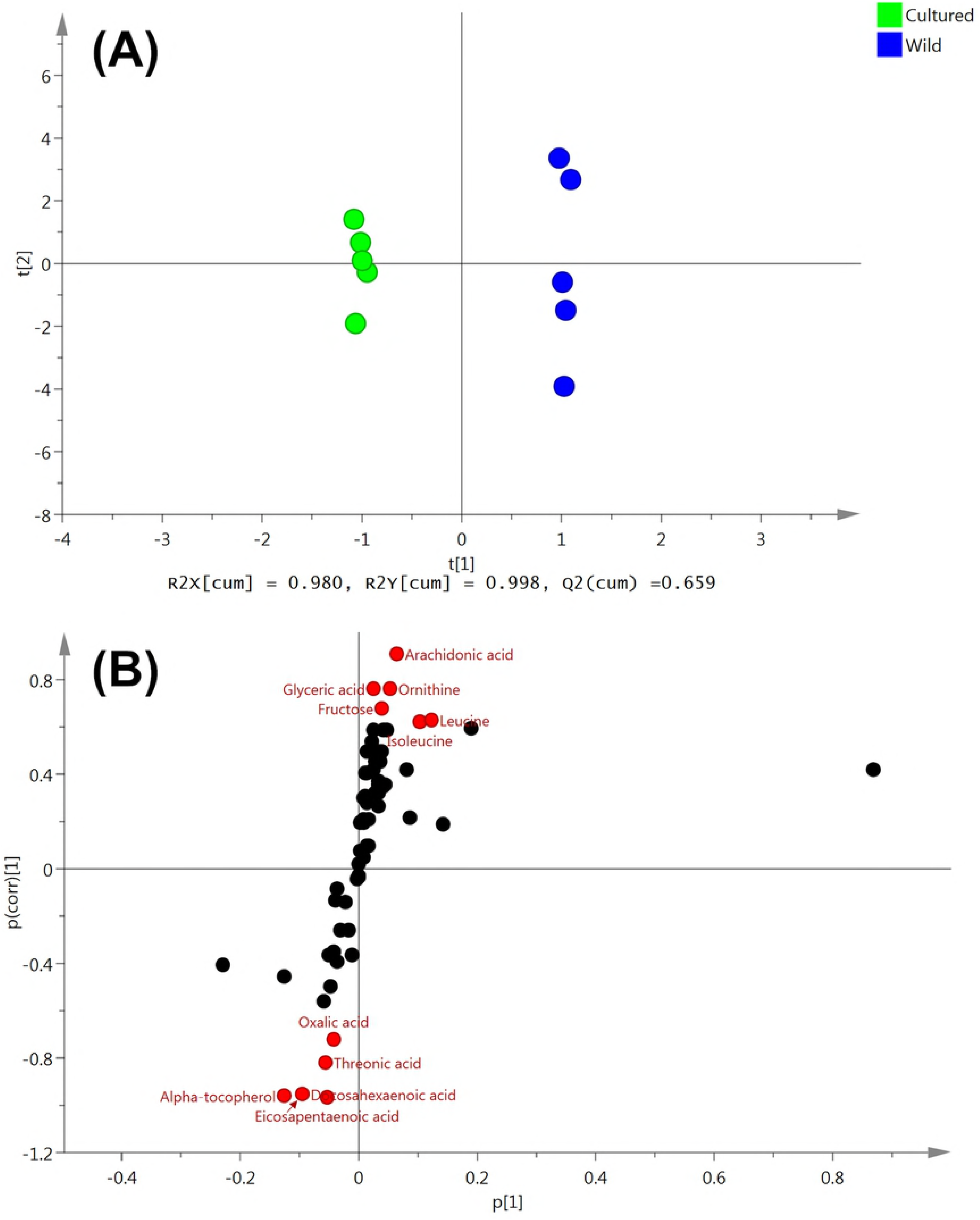
OPLS-DA of serum metabolites obtained from cultured and wild eels. (A) OPLS-DA of serum metabolites obtained from cultured and wild eels. (B) S-plot generated from OPLS-DA.

In the liver, we identified 8 metabolites as candidate biomarkers; wild eels showed high levels of 1, whereas cultured eels showed high levels of 7 (Table 4). In cultured eels, sugars (glucose-6-phosphate and maltose) showed a about 5-fold increase, which may have important implications for energy metabolism disorders.

**Table 4.** Fold change of biomarker candidates between cultured and wild eel liver * and ** denotes significant difference at *P < 0.05* and *P < 0.01*, respectively.

| Metabolites | Fold change<br>(Cultured/Wild) | *P*-value |
| --- | --- | --- |
| Arachidonic acid | 0.22 | * |
| Myoinositol-1-phosphate | 1.34 |  |
| Docosahexaenoic acid | 1.90 | ** |
| Citric acid | 1.91 | ** |
| Eicosapentaenoic acid | 1.97 | ** |
| Glucose-6-phosphate | 4.96 | ** |
| Maltose | 5.66 |  |
| Alpha-tocopherol | 9.45 | * |

In the ovaries, 11 metabolites were identified as candidate biomarkers; wild eels showed high levels of 6, whereas cultured eels showed high levels of 5 (Table 5). Pantothenic acid (vitamin B5) was uniquely detected only in the ovaries, and increased by 1.8-fold in cultured eels. Unfortunately, one sample of organic phase extract from cultured eel ovary was not correctly analyzed by GC/MS, therefore, this sample was omitted from analysis.

**Table 5.** Fold change of biomarker candidates between cultured and wild eel ovary. and ** denotes significant difference at *P < 0.05* and *P < 0.01*, respectively.

| Metabolites | Fold change<br>(Cultured/Wild) | *P*-value |
| --- | --- | --- |
| Arachidonic acid | 0.37 | ** |
| Pyroglutamic acid | 0.43 |  |
| Elaidic acid | 0.53 | * |
| Alpha-ketoglutaric acid | 0.57 | * |
| Oleic acid | 0.74 |  |
| Beta-alanine | 0.76 | * |
| Docosahexaenoic acid | 1.50 | * |
| Panthothenic acid | 1.81 | ** |
| Oxalic acid | 1.84 | * |
| Eicosapentaenoic acid | 2.03 | ** |
| Alpha-tocopherol | 5.27 | ** |

\* and \*\* denotes significant difference at $P < 0.05$ and $P < 0.01$ , respectively.

In the serum samples, we identified 11 metabolites as candidate biomarkers; wild eels show high levels of 6, whereas cultured eels showed high levels of 5 (Table 6). Along with eicosapentaenoic acid (EPA), docosahexaenoic acid (DHA) showed a significant increase in cultured eels, while arachidonic acid (ARA) was found in high levels in wild eels. Threonic acid and oxalic acid, which are metabolites of ascorbate, increased in cultured eels.

**Table 6.** Fold change of biomarker candidates between cultured and wild eel serum. and ** denotes significant difference at *P < 0.05* and *P < 0.01*, respectively.

| Metabolites | Fold change<br>(Cultured/Wild) | *P-value* |
| --- | --- | --- |
| Arachidonic acid | 0.14 | ** |
| Fructose | 0.30 |  |
| Ornithine | 0.45 | * |
| Isoleucine | 0.53 |  |
| Leucine | 0.55 |  |
| Glyceric acid | 0.58 | * |
| Oxalic acid | 1.58 | * |
| Threonic acid | 2.64 | ** |
| Docosahexaenoic acid | 2.81 | ** |
| Eicosapentaenoic acid | 2.84 | ** |
| Alpha-tocopherol | 3.18 | ** |

It is noteworthy that EPA, DHA, and α-tocopherol (lipophilic vitamin E) increased in all three tissue samples of cultured eel, while ARA increased in all three tissue samples of wild eels. The DHA/EPA/ARA ratios calculated by normalized average peak area (peak areas of each fatty acids/peak area of internal standard/sample weight-dry) by GC/MS analyses in cultured eels and wild eels were 10.1/3.2/1.0 and 1.1/0.3/1.0, respectively in the liver; 4.0/1.9/1.0 and 1.0/0.4/1.0, respectively in the ovaries; and 19.5/6.5/1.0 and 1.1/0.3/1.0, respectively, in the serum.

### TUNEL assay

We examined 5 eel ovarian samples from each eel group. We detected TUNEL-positive cells only in cultured eel ovaries (Fig 7) in interstitial cells. The oocytes in both cultured and wild eels were mainly in the late-cortical alveoli to early-vitellogenic developmental stage.

**Fig 7.**
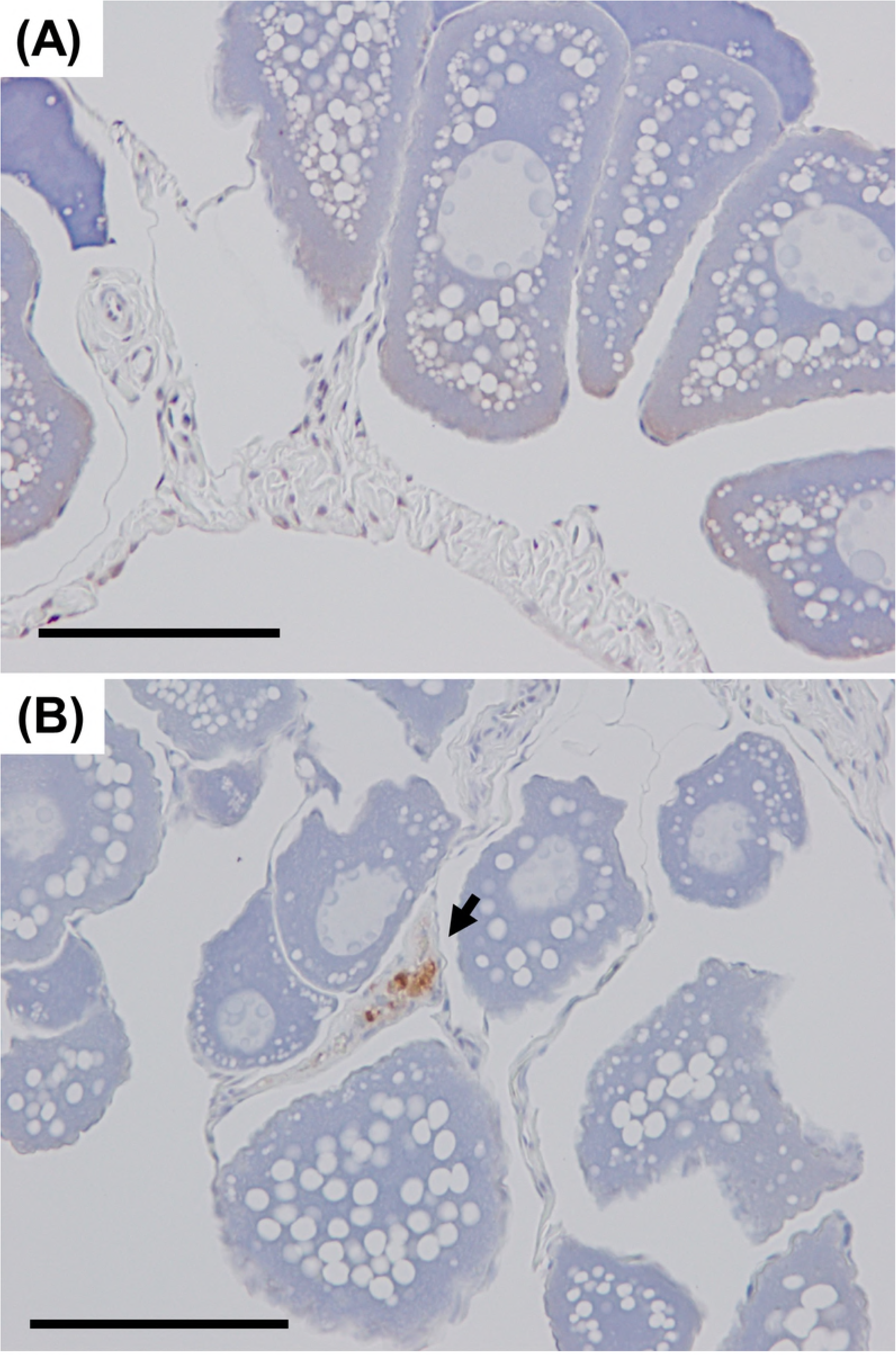
TUNEL assay in eel ovary samples. (A) wild eel, (B) cultured eel. Arrowhead: TUNEL positive cells. Bars, = 100 μm.

## Discussion

In this study, we clearly showed the differences in nutritional status just before the beginning of artificial induction of oocyte maturation between cultured and wild eels using trans-omics analysis. We also demonstrated the influence of these differences on the eel’s hormonal status. Results of this comprehensive study suggested that the main cause of ovulation problem, frequently found in cultured eels, is nutritional status.

Transcriptomic analysis revealed that the transcription level of *gh* in the cultured eel brain is markedly lower than in wild eels. In teleosts, growth hormone (GH) affects several aspects of behavior, including appetite, foraging, aggression, and predator avoidance [16–18], and reportedly affects reproduction, including gonad development [19, 20], and steroidogenesis promotion [21]. In human, GH plays major role in the control of ovarian function in conjunction with the gonadotropins, including puberty, gametogenesis, steroidogenesis, and ovulation [22–24]; however, no data are available on the effect of GH on ovulation in teleosts. Interestingly, studies on the Asian catfish have demonstrated that GH injection in the morning in the early recrudescence phase induce an appreciable increase in ovarian weight and the GSI [25]. In our study, the ovarian weight and GSI of cultured eels were found to lower than those of wild eels. Our RNA-seq-based transcriptomic analysis data indicate recognizable transcription of *gh receptor 1, gh receptor 2*, and *gh-releasing hormone receptor*. However, the transcription levels of gonadotropin receptors were very low. Studies have reported changes in gonadotropin receptors in the ovaries during hormone-induced oocyte maturation in the Japanese eel [26]. In the late vitellogenic stage, transcription levels of *luteinizing hormone receptor* in the ovaries increase. Taken together, the results indicated that compared to low *lh* levels, low *gh* levels mainly influence lower ovarian weight in cultured eels.

This study also demonstrated that *shbg* transcription levels in the liver is low in cultured eels. In mammals, GH pulsatility or thyroid hormone is considered to enhance SHBG transcription *via* hepatocyte nuclear factor-4alpha (HNF4a) [27, 28]. These data suggest that not only *gh* transcription but also GH secretion is lower in cultured eels compared to wild eels, although no significant difference was observed in the transcription levels of *insulin-like growth factor ?* (*igf1*) in the liver in the two eel groups. Initially, hepatic IGF1 expression was thought to be fully dependent on GH. However, studies have reported that the total serum IGF1 concentration does not significantly differ between the human obese and control groups, despite GH hyposecretion in obese people [29]. In addition, glucose and insulin can also promote IGF1 expression in fetal rat hepatocytes using *in vitro* [30]. These data indicate GH and IGF1 levels are not always correlated. In addition, Igf1and Igf2 are expressed in granulosa cells of fish oocytes [31, 32], and it is hypothesized that in the red seabream, Igf1 increases the responsiveness to luteinizing hormone (LH) in E2 production through an increase in LH receptor gene expression [33]. Thus, further studies are needed in order to clarify the molecular mechanisms of diminished GH secretion on final maturation and ovulation via Igf regulation in the Japanese eel. Further, we also think that the cultured eels in our study were not obese because their condition factor was lower than the that of wild eels. The condition factor is used to evaluate the fatness and well-being of fish. Therefore, signs and symptoms in cultured eels we observed in our study, such as low *gh* levels, low *shbg* levels, and ovulation problem are associated with human non-obese PCOS [23, 34–36].

Transcriptomic analysis also demonstrated that apoptosis-associated genes such as *small nuclear ribonucleoprotein F, tumor necrosis factor receptor superfamily member 19, cytochrome c*, and *granzyme K*, are highly expressed in cultured eel ovaries and that *cytochrome c* is highly expressed in cultured eel liver [37, 38]. In this study, TUNEL assay demonstrated apoptotic cells only in cultured eel ovaries. These data indicate that the caspase-independent pathway of programmed cell death occurs in cultured eel ovaries, and many reports have shown that GH suppresses apoptosis [39–41]. Taken together, the results indicated that impaired GH secretion is likely to induce apoptosis in cultured eel ovaries and probably in the liver, too.

Currently, the main cause of PCOS is thought to be insulin resistance, which, in turn, is related to hyperinsulinemia. In the past, hyperinsulinemia was known as metabolic syndrome [42]. A high-fat and/or high-carbohydrate diet is considered the major cause of insulin resistance [43, 44]. In our study, metabolomic analysis using GC/MS showed intense accumulation of glucose-6-phosphate and maltose in the cultured eel liver, while no significant difference in the serum glucose levels between wild and cultured eels.

Commercial diet for the Japanese eel contains 23%–24% potato starch as a binder (See Table 1). In vertebrate, glucose is phosphorylated to glucose-6-phosphate and stored in the liver. Homogenization of eel pancreatic segments showed α-amylase activity [45], indicating that starch must be digested to glucose or maltose in the eel’s digestive tract. This suggests that accumulated glucose-6-phosphate and maltose in the cultured eel liver are mainly derived from starch present in the commercial diet. In teleosts, unlike in mammals, amino acids play a more significant role in insulin secretion than does glucose [46, 47], and teleosts have been generally considered glucose intolerant [48]. However, glucose stimulates insulin and somatostatin-14 production, which suppress GH secretion, and inhibits glucagon secretion in angler fish [49]. In addition, when arginine and glucose were simultaneously injected into the European eel, insulin secretion stimulation was greater than after injection of either factor alone [50]. The high insulin level probably suppresses glucose release and promotes glucose storage. These results indicated that cultured eels have hyperinsulinemia, in addition to significantly high serum triglyceride levels. In future, serum insulin levels in cultured eels should be measured.

Our results suggested that the significantly low transcription level of *shbg* in cultured eels compare to wild eels is reflected by lower transcription levels of GH. Shbg characterization in the Japanese eel has been reported earlier [51]. In the Japanese eel, Shbg showed high affinity to E2, androstenediol and T. Shbg in other fish species such as salmonids, goldfish, and zebrafish is also well characterized [52–54]. In trout, Shbg-bound E2 has the highest affinity, followed by T. In zebrafish, Shbg-bound T and androstenedione have the highest affinity, followed by E2 and 5α-DHT. Although the *shbg* transcription levels in cultured eels is lower than wild eels, there is no significant difference in serum steroid hormone levels between the two eel groups. However, circulating T levels tend to be relatively lower, and the hepatic *akr1d1* mRNA level is significantly higher in cultured eels. Akr1d1, the steroid 5β-reductase, is responsible for conversion of T to 5β-DHT [55], and low SHBG level is associated with an increasing free T level. Taken together the results suggested that in cultured eels, increased free T undergoes clearance in the liver by sequential reduction by Akr1d1. Unfortunately, we were unable to measure the serum level of 5β-DHT because of a lack of an appropriate assay. Further, serum 11-KT, a major androgen in this species, was slightly but not significantly higher in wild eels. The transcription level of *11β hydroxysteroid dehydrogenase short form* (*11β-hsdsf*), which catalyzes 11β-hydroxytestosterone to 11-KT and cortisol to cortisone [56], was also higher in the wild eel liver compared to cultured eels. Therefore, high serum 11-KT levels in wild eels may be associated with higher *11β-hsdsf* transcription in the liver.

In humans, the low Shbg level is also associated with hyperandrogenism. Therefore, we examined whether hyperandrogenic symptoms are also observed in cultured eels. Initially, marker genes selected on the basis of transcriptomic analysis result between wild female and wild male brains showed sex-dependent expression. As mentioned before, *sl* and *cyp19a1* were selected as female highly expressed genes, and *jumonji* was selected as a male highly expressed gene. Strangely, in cultured eels, we could not find hyperandrogenic symptoms due to low *shbg* levels. Several symptoms such as lower *gh* and *shbg* transcription levels, ovulation problem in cultured eels are associated with human PCOS. However, hyperandrogenism were not observed in cultured eels. Therefore, it is thought that the condition of cultured eels is a little different from human PCOS. Furthermore, E2 or 11-KT serum levels increased toward the final maturation stage in eel [57]. Therefore, further studies are required to examine the effects of lower Shbg levels on steroid hormone profiles during artificial oocyte maturation.

Although we could not find hyperandrogenic symptoms in cultured eels there were significant sex differences in gene expression in brains. Naturally, the expression level of *cyp19a1*, also called *aromatase*, is higher in females but it is unclear why the *sl* levels is higher in females. Several studies have reported that in the Japanese eel, the numbers of anti-Sl-positive cells in pituitary gradually increase during artificial oocyte maturation until the late vitellogenic stage and then decrease at the migratory nucleus stage [58]. Also, relative mRNA levels of *sl* in the pituitary of blue gourami (*Trichogaster trichopterus*) gradually decreased during oocyte maturation [59]. These data suggest that Sl plays a role in oocyte maturation. Many JUMONJI family proteins have domains, such as ARID, jmjC, and jmjN, involved in DNA binding, chromatin binding, and transcription, suggesting that JUMONJI family proteins regulate transcription or chromatin function or both [60]. In this study, transcriptomic analysis showed that mRNA expression levels of the four *jumonji* family genes are higher in the male eel brain. All four genes contain ARID, jmjC and/or jmjN domains (data not shown). Interestingly, studies have demonstrated that JMDM2A, a jmjC-containing histone H3 lysine 9 (H3K9) demethylase, interacts directly with the androgen receptor (AR) and is employed in a hormone-dependent manner in AR target genes in order to mediate transcription activation using human cell lines [61]. These data suggest that *jumonji* family genes may contribute to sex differences in the Japanese eel brain.

Trans-omics analysis provides us with “the big picture", and also gives us other insights. In this study, we found α-tocopherol accumulation in cultured eel liver, ovary, and serum. Other studies have reported that excess α-tocopherol induces tocopherol radicals and oxidizes ascorbic acid to reduce oxidized tocopherol [62]. Oxidized ascorbic acid, also called dehydroascorbic acid, is degraded to threonic acid and oxalic acid [63]. In cultured eels, metabolomic analysis showed high oxalic acid levels in the ovaries and high oxalic acid and threonic acid levels in the serum. In addition, ascorbic acid and dehydroascorbic acid levels in ovaries of cultured eels were a little lower compared to wild eels (data not shown). These data indicate that cultured eels consume excessive α-tocopherol. Metabolomic analysis also revealed that the DHA/EPA/ARA ratio in all examined samples of cultured eels are quite different from that in wild eels. In our study, cultured eels were fed diet supplemented with cod liver oil (Table 2). Cod liver oil contains much DHA and EPA compare to ARA [64].

Therefore, it is indicated that dietary DHA/EPA/ARA ratio affected tissue DHA/EPA/ARA ratio in cultured eels. Lower ARA level in eggs from cultured Japanese eel than wild eel has also been reported [65]. In addition, several reports have indicated the DHA/EPA/ARA ratio in ovaries or eggs strongly influences egg and larval quality in fish [66–69]. ARA is the precursor of biologically active eicosanoids, such as prostaglandins, which are involved in many aspects of reproduction including reproductive behavior and ovulation in teleosts [70, 71]. EPA and ARA compete for the enzymatic pathways involved in synthesis of eicosanoids; therefore, further studies are required in order to examine a dietary DHA/EPA/ARA ratio optimal for reproductive performance.

## Conclusion

In conclusion, trans-omics analysis indicated that ovulation problem, found frequently in cultured eels was due to a high-carbohydrate diet and/or unbalanced DHA/EPA/ARA ratios in diet. Nutritional interactions with reproductive performance have been reported in domesticated animals [72-74]. It has been reported that cultured broodstock showed poor reproductive performance compared to wild fish in several fish species [75-77]. The commercial diet for fish is generally designed for weight gain and/or fleshing out, regardless of health, and optimized diet for broodstock culture is not commercially available. Therefore, dietary effects on the reproductive performance of fish should be examined more deeply in the future. Our study suggested that effects of high-carbohydrate diet on GH secretion, effects of impaired GH secretion on responsiveness to gonadotropins, effects of lower Shbg levels on oocyte maturation, and optimal dietary DHA/EPA/ARA ratios, are future experimental directions for improvement of seed production efficiency. The comprehensive data obtained from our study also demonstrate the usefulness of omics approaches in understanding the functionality of diet components in aquaculture.

## Acknowledgement

The authors declare no conflicts of interest associated with this manuscript. We thank Ei-ichi Yamamoto for his attentive care for eels, and Dr. Katsutoshi Ito and Dr. Mana Ito for their technical advices.

## Author contribution

Conceptualization: MH.

Data curation: MH.

Formal analysis: MH. MM. TH.

Investigation: MH. HI.

Methodology: MH. MM. TH.

Resources: MH. HI.

Validation: MH. MM. TH.

Visualization: MH. TH.

Writing – original draft: MH.

## Supporting Information

Supplemental table S1. Primer and probe sequences for RT-qPCR. primer and probe sequences for RT-qPCR in this research

Supplemental table S2. Comparison of gene expression in liver. List of the significant differentially expressed genes in wild and cultured eel liver obtained from our RNA-seq data

Supplemental table S3. Comparison of gene expression in ovary. List of significant differentially expressed genes in wild and cultured eel ovary

Supplemental table S4. Comparison of gene expression in brain between wild eel female and cultured eel female. List of significant differentially expressed genes in wild female brain and wild female brain

Supplemental table S5. Comparison of gene expression in brain between wild eel female and wild male eel. List of significant differentially expressed genes in wild female brain and wild male brain

